## Supplemental Figures for "Molecular, Cellular, and Developmental Organization of the Mouse Vomeronasal organ at Single Cell Resolution"

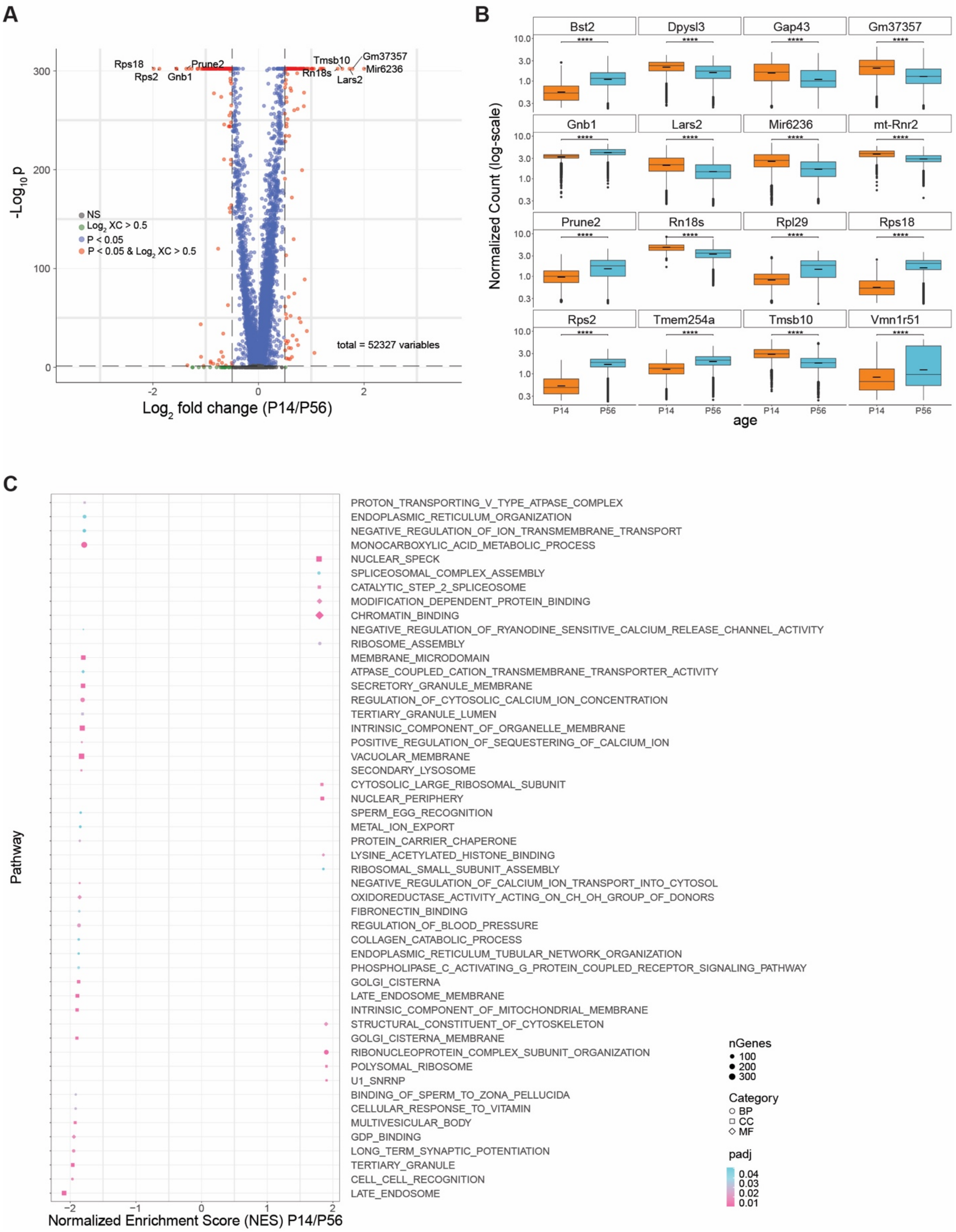
A
Rps18
Rps2
Gnb1
Prune2
Tmsb10
Gm37357
Mir6236
Rn18s
Lars2
NS
Log2 XC > 0.5
P < 0.05
P < 0.05 & Log2 XC > 0.5
total = 52327 variables
-Log10 p
Log2 fold change (P14/P56)
B
Bst2
Dpysl3
Gap43
Gm37357
Gnb1
Lars2
Mir6236
mt-Rnr2
Prune2
Rn18s
Rpl29
Rps18
Rps2
Tmem254a
Tmsb10
Vmn1r51
Normalized Count (log-scale)
P14
P56
age
C
PROTON_TRANSPORTING_V_TYPE_ATPASE_COMPLEX
ENDOPLASMIC_RETICULUM_ORGANIZATION
NEGATIVE_REGULATION_OF_ION_TRANSMEMBRANE_TRANSPORT
MONOCARBOXYLIC_ACID_METABOLIC_PROCESS
NUCLEAR_SPECK
SPLICEOSOMAL_COMPLEX_ASSEMBLY
CATALYTIC_STEP_2_SPLICEOSOME
MODIFICATION_DEPENDENT_PROTEIN_BINDING
CHROMATIN_BINDING
NEGATIVE_REGULATION_OF_RYANODINE_SENSITIVE_CALCIUM_RELEASE_CHANNEL_ACTIVITY
RIBOSOME_ASSEMBLY
MEMBRANE_MICRODOMAIN
ATPASE_COUPLED_CATION_TRANSMEMBRANE_TRANSPORTER_ACTIVITY
SECRETORY_GRANULE_MEMBRANE
REGULATION_OF_CYTOSOLIC_CALCIUM_ION_CONCENTRATION
TERTIARY_GRANULE_LUMEN
INTRINSIC_COMPONENT_OF_ORGANELLE_MEMBRANE
POSITIVE_REGULATION_OF_SEQUESTERING_OF_CALCIUM_ION
VACUOLAR_MEMBRANE
SECONDARY_LYSOSOME
CYTOSOLIC_LARGE_RIBOSOMAL_SUBUNIT
NUCLEAR_PERIPHERY
SPERM_EGG_RECOGNITION
METAL_ION_EXPORT
PROTEIN_CARRIER_CHAPERONE
LYSINE_ACETYLATED_HISTONE_BINDING
RIBOSOMAL_SMALL_SUBUNIT_ASSEMBLY
NEGATIVE_REGULATION_OF_CALCIUM_ION_TRANSPORT_INTO_CYTOSOL
OXIDOREDUCTASE_ACTIVITY_ACTING_ON_CH_OH_GROUP_OF_DONORS
FIBRONECTIN_BINDING
REGULATION_OF_BLOOD_PRESSURE
COLLAGEN_CATABOLIC_PROCESS
ENDOPLASMIC_RETICULUM_TUBULAR_NETWORK_ORGANIZATION
PHOSPHOLIPASE_C_ACTIVATING_G_PROTEIN_COUPLED_RECEPTOR_SIGNALING_PATHWAY
GOLGI_CISTERNA
LATE_ENDOSOME_MEMBRANE
INTRINSIC_COMPONENT_OF_MITOCHONDRIAL_MEMBRANE
STRUCTURAL_CONSTITUENT_OF_CYTOSKELETON
GOLGI_CISTERNA_MEMBRANE
RIBONUCLEOPROTEIN_COMPLEX_SUBUNIT_ORGANIZATION
POLYSOMAL_RIBOSOME
U1_SNRNP
BINDING_OF_SPERM_TO_ZONA_PELLUCIDA
CELLULAR_RESPONSE_TO_VITAMIN
MULTIVESICULAR_BODY
GDP_BINDING
LONG_TERM_SYNAPTIC_POTENTIATION
TERTIARY_GRANULE
CELL_CELL_RECOGNITION
LATE_ENDOSOME
Pathway
Normalized Enrichment Score (NES) P14/P56
nGenes
100
200
300
Category
BP
CC
MF
padj
0.04
0.03
0.02
0.01

Fig. S1. Age differences in gene expression.

**A** Volcano plot of gene expression differences between P14 and P56. **B**. 16 significantly differentially expressed genes with largest positive or negative $\log_2$-fold-change values. **C**. 50 significantly enriched GO terms with largest positive or negative $\log_2$-fold-change values.

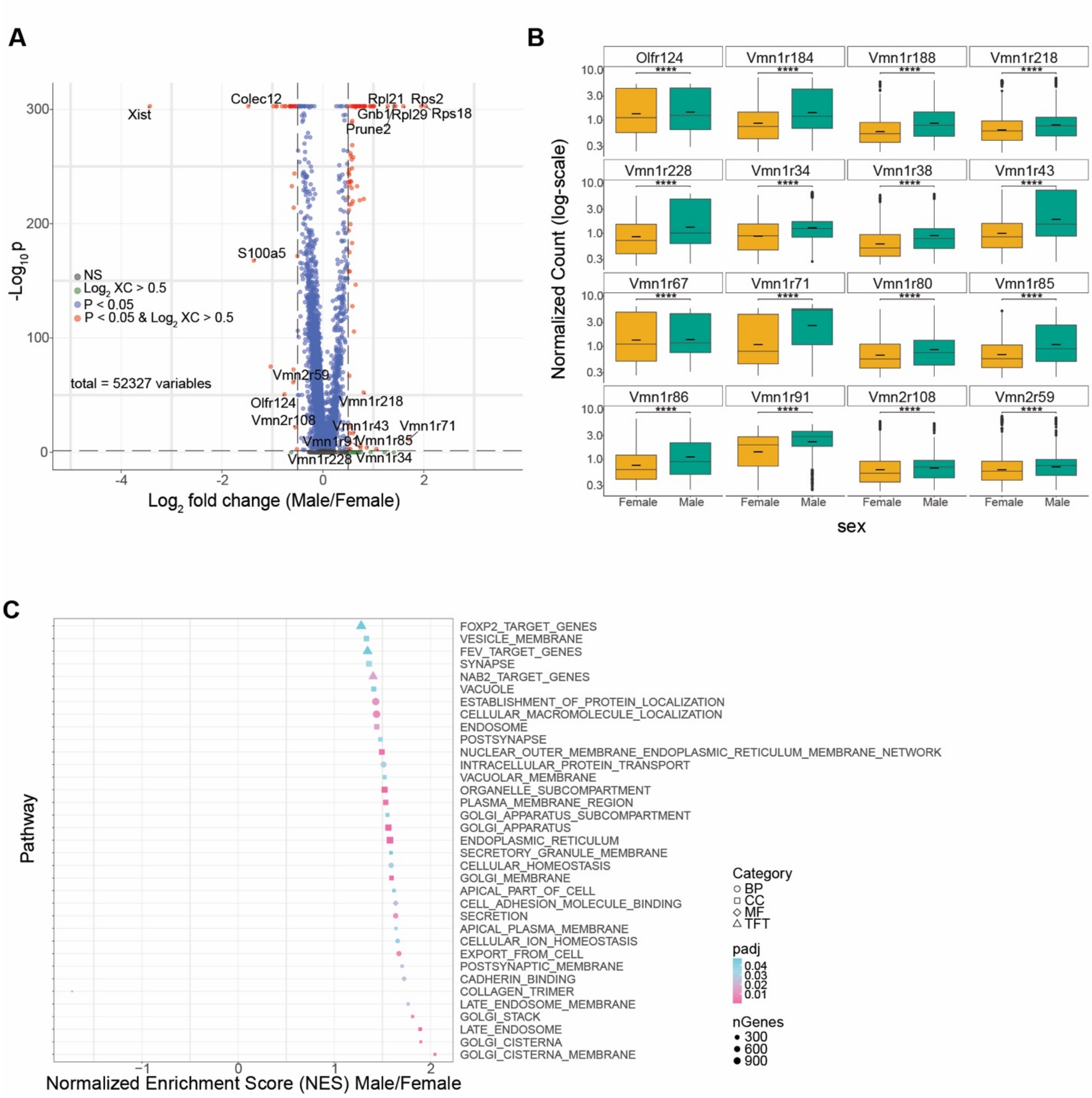


Fig. S2. Sex differences in gene expression.
**A** Volcano plot of gene expression differences between male and female mice. **B**. 16 significantly differentially expressed chemosensory receptors with largest positive or negative $\log_2$-fold-change values. **C**. 35 significantly enriched GO terms.

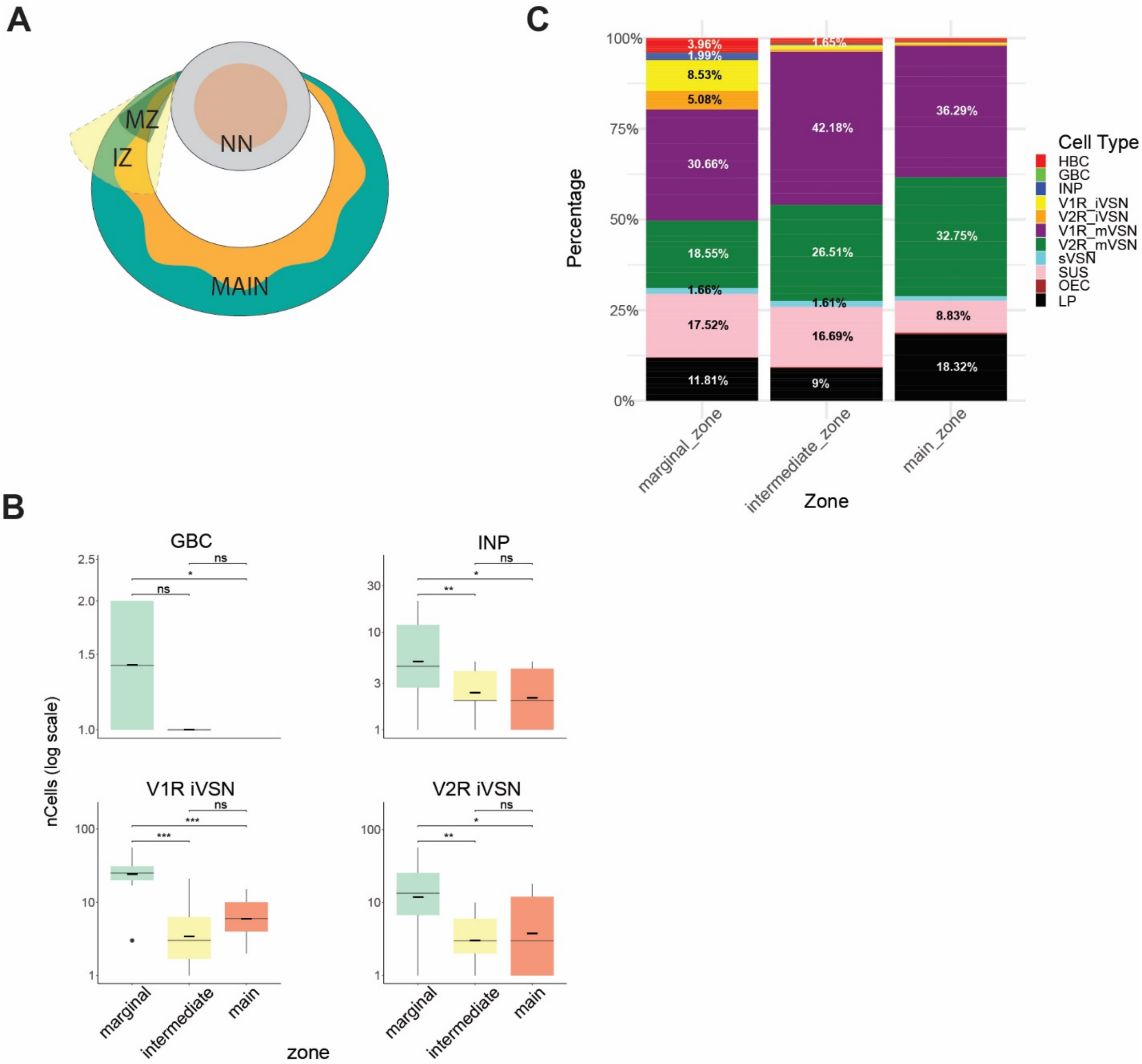


Fig. S3. Zonal distribution of cell types in the VNO neuroepithelia.
**A.** Schematic indicating quantification of cells in the marginal, intermediate, and main zones.
**B**. Stacked bar-plot of cell-type proportions by VNO zone. **C**. Box plots of GBC, INP, and immature VSN cell counts by zone, across 13 slides.

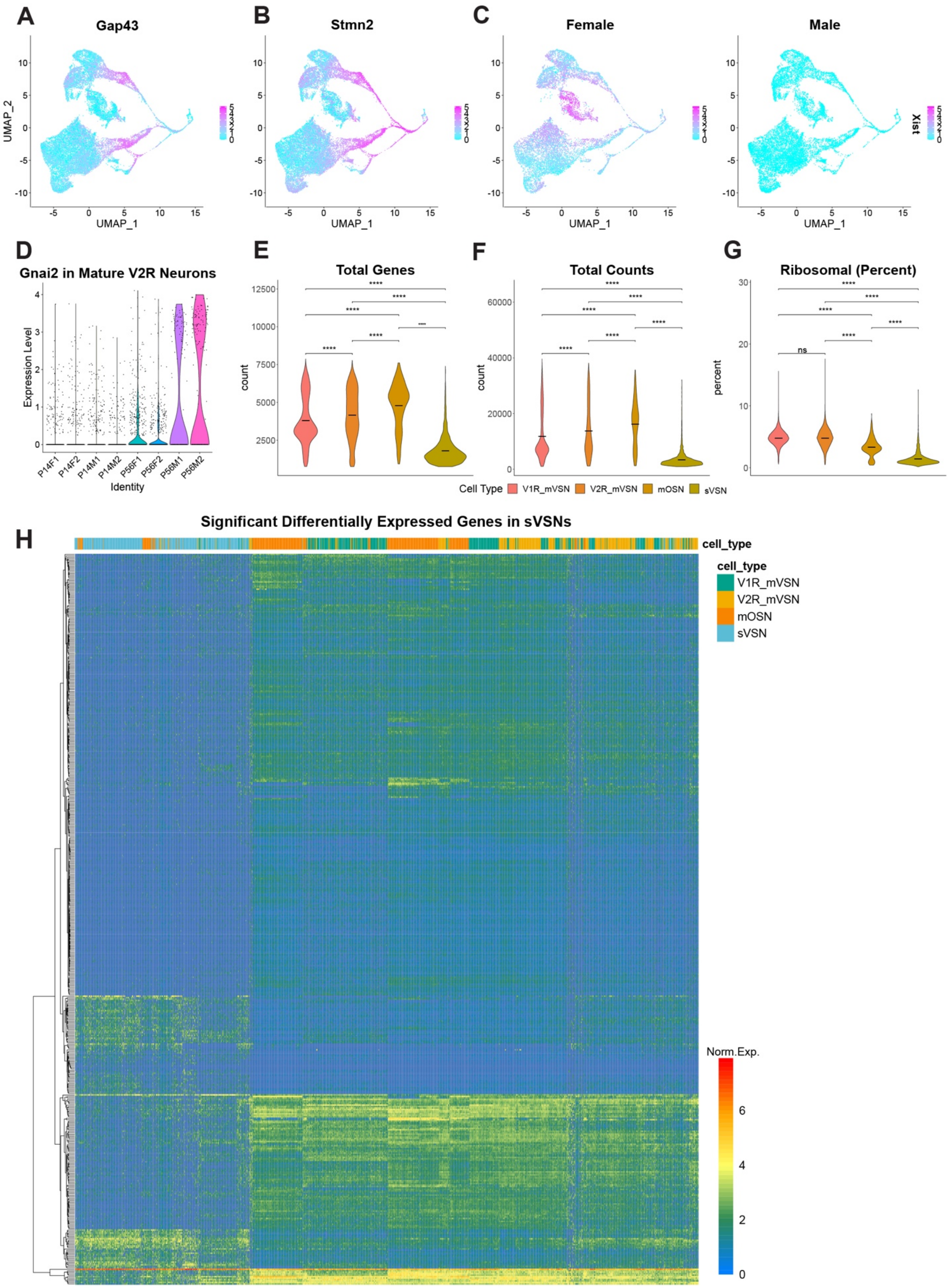
A
Gap43
B
Stmn2
C
Female
Male
Xist
UMAP_1
UMAP_2
D
Gnai2 in Mature V2R Neurons
Expression Level
Identity
P14F1
P14F2
P14M1
P14M2
P56F1
P56F2
P56M1
P56M2
E
Total Genes
F
Total Counts
G
Ribosomal (Percent)
count
percent
ns
Cell Type
V1R_mVSN
V2R_mVSN
mOSN
sVSN
H
Significant Differentially Expressed Genes in sVSNs
cell_type
Norm.Exp.

Fig. S4. Neuronal lineage features and differentially expressed genes in sVSNs.

**A.** and **B**. Normalized Gap43 and Stmn2 expression in the neuronal lineage. **C**. Normalized Xist expression in the neuronal lineage, split by sex. **D**. Violin plot of normalized Gnai2 expression in mature V2R neurons, split by sample. **E-G**. Total number of genes, counts, and percent ribosomal gene expression detected in mature neurons, split by cell type. **H**. Heatmap of 503 significant ($p_{adj} \leq 0.001$, fold-change $\geq 1.5$) differentially expressed genes in sVSNs.

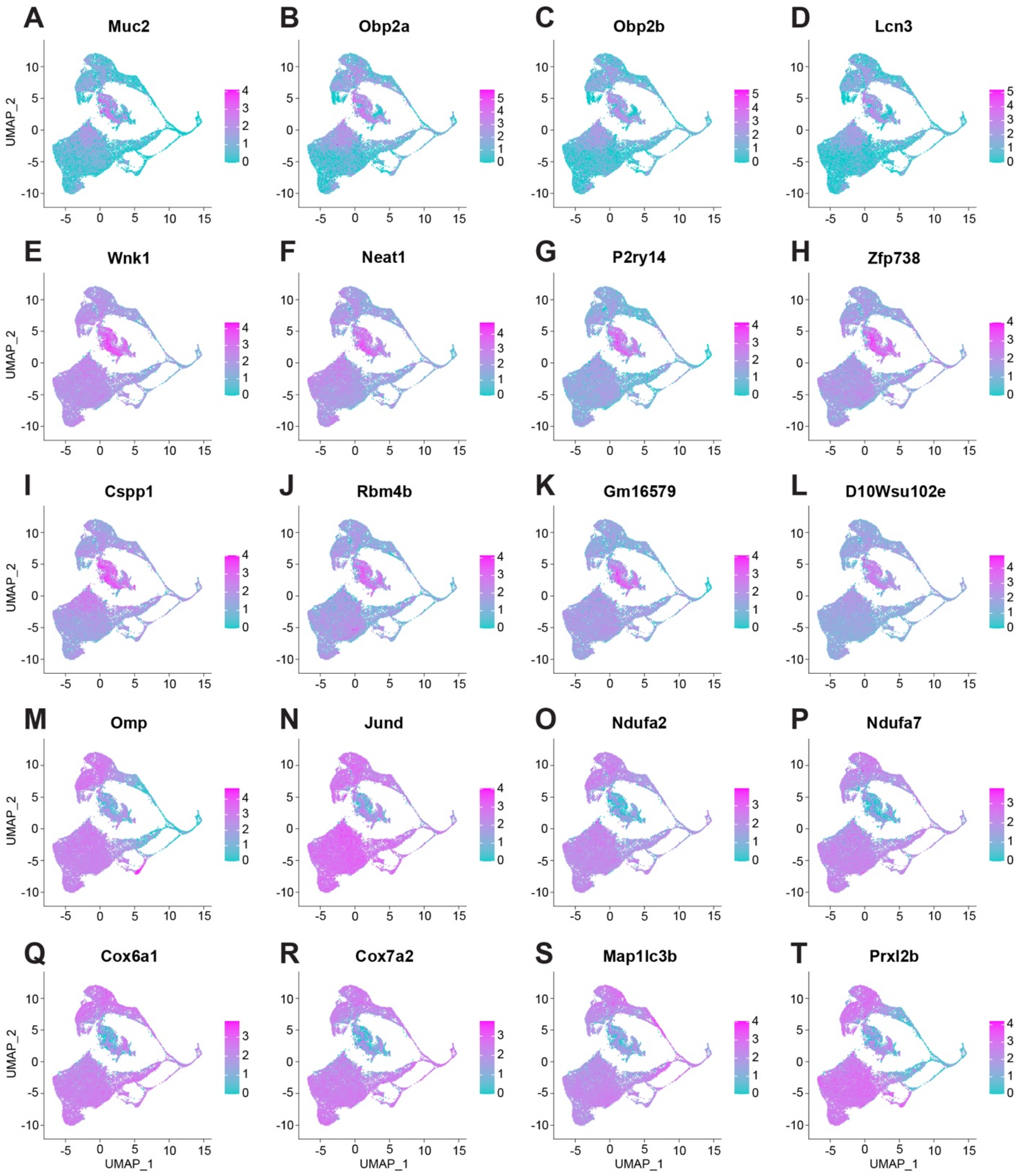


Fig. S5. Significantly regulated genes in sVSNs.

**A-D.** Feature plots showing odor binding protein gene expression (Muc2, Obp2a, Obp2b, and Lcn3). **E-L**. Top upregulated genes in sVSNs. **M-T**. Top downregulated genes in sVSNs.

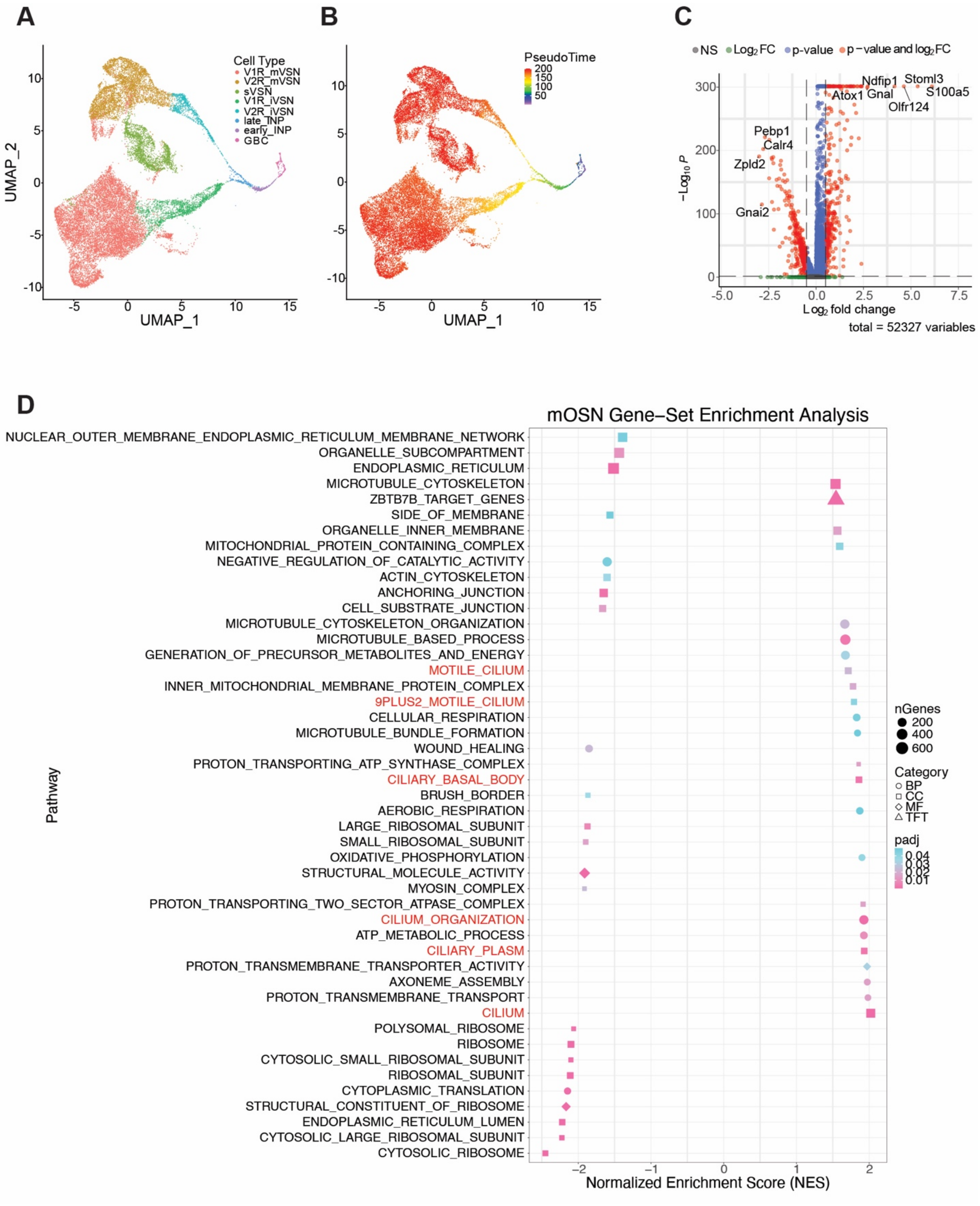
A
B
C
D
Cell Type
V1R_mVSN
V2R_mVSN
sVSN
V1R_iVSN
V2R_iVSN
late_INP
early_INP
GBC
UMAP_1
UMAP_2
PseudoTime
NS
Log2 FC
p-value
p-value and log2 FC
Ndfip1
Stoml3
S100a5
Atox1
Gnal
Olfr124
Pebp1
Calr4
Zpld2
Gnai2
-Log10 P
Log2 fold change
total = 52327 variables
mOSN Gene-Set Enrichment Analysis
NUCLEAR_OUTER_MEMBRANE_ENDOPLASMIC_RETICULUM_MEMBRANE_NETWORK
ORGANELLE_SUBCOMPARTMENT
ENDOPLASMIC_RETICULUM
MICROTUBULE_CYTOSKELETON
ZBTB7B_TARGET_GENES
SIDE_OF_MEMBRANE
ORGANELLE_INNER_MEMBRANE
MITOCHONDRIAL_PROTEIN_CONTAINING_COMPLEX
NEGATIVE_REGULATION_OF_CATALYTIC_ACTIVITY
ACTIN_CYTOSKELETON
ANCHORING_JUNCTION
CELL_SUBSTRATE_JUNCTION
MICROTUBULE_CYTOSKELETON_ORGANIZATION
MICROTUBULE_BASED_PROCESS
GENERATION_OF_PRECURSOR_METABOLITES_AND_ENERGY
MOTILE_CILIUM
INNER_MITOCHONDRIAL_MEMBRANE_PROTEIN_COMPLEX
9PLUS2_MOTILE_CILIUM
CELLULAR_RESPIRATION
MICROTUBULE_BUNDLE_FORMATION
WOUND_HEALING
PROTON_TRANSPORTING_ATP_SYNTHASE_COMPLEX
CILIARY_BASAL_BODY
BRUSH_BORDER
AEROBIC_RESPIRATION
LARGE_RIBOSOMAL_SUBUNIT
SMALL_RIBOSOMAL_SUBUNIT
OXIDATIVE_PHOSPHORYLATION
STRUCTURAL_MOLECULE_ACTIVITY
MYOSIN_COMPLEX
PROTON_TRANSPORTING_TWO_SECTOR_ATPASE_COMPLEX
CILIUM_ORGANIZATION
ATP_METABOLIC_PROCESS
CILIARY_PLASM
PROTON_TRANSMEMBRANE_TRANSPORTER_ACTIVITY
AXONEME_ASSEMBLY
PROTON_TRANSMEMBRANE_TRANSPORT
CILIUM
POLYSOMAL_RIBOSOME
RIBOSOME
CYTOSOLIC_SMALL_RIBOSOMAL_SUBUNIT
RIBOSOMAL_SUBUNIT
CYTOPLASMIC_TRANSLATION
STRUCTURAL_CONSTITUENT_OF_RIBOSOME
ENDOPLASMIC_RETICULUM_LUMEN
CYTOSOLIC_LARGE_RIBOSOMAL_SUBUNIT
CYTOSOLIC_RIBOSOME
Pathway
Normalized Enrichment Score (NES)
nGenes
Category
BP
CC
MF
TFT
padj

Fig. S6. sVSN Pseudotime, OSN markers and GO Terms.
**A.** VNO Neuronal lineage cell-types in UMAP space. **B**. V1R, V2R, and sVSN lineage pseudotimes in UMAP space. **C**. Volcano plot of mOSN gene expression versus all other cells from the neuronal lineage. **D.** 45 significant GO terms enriched in OSNs versus all other cells.

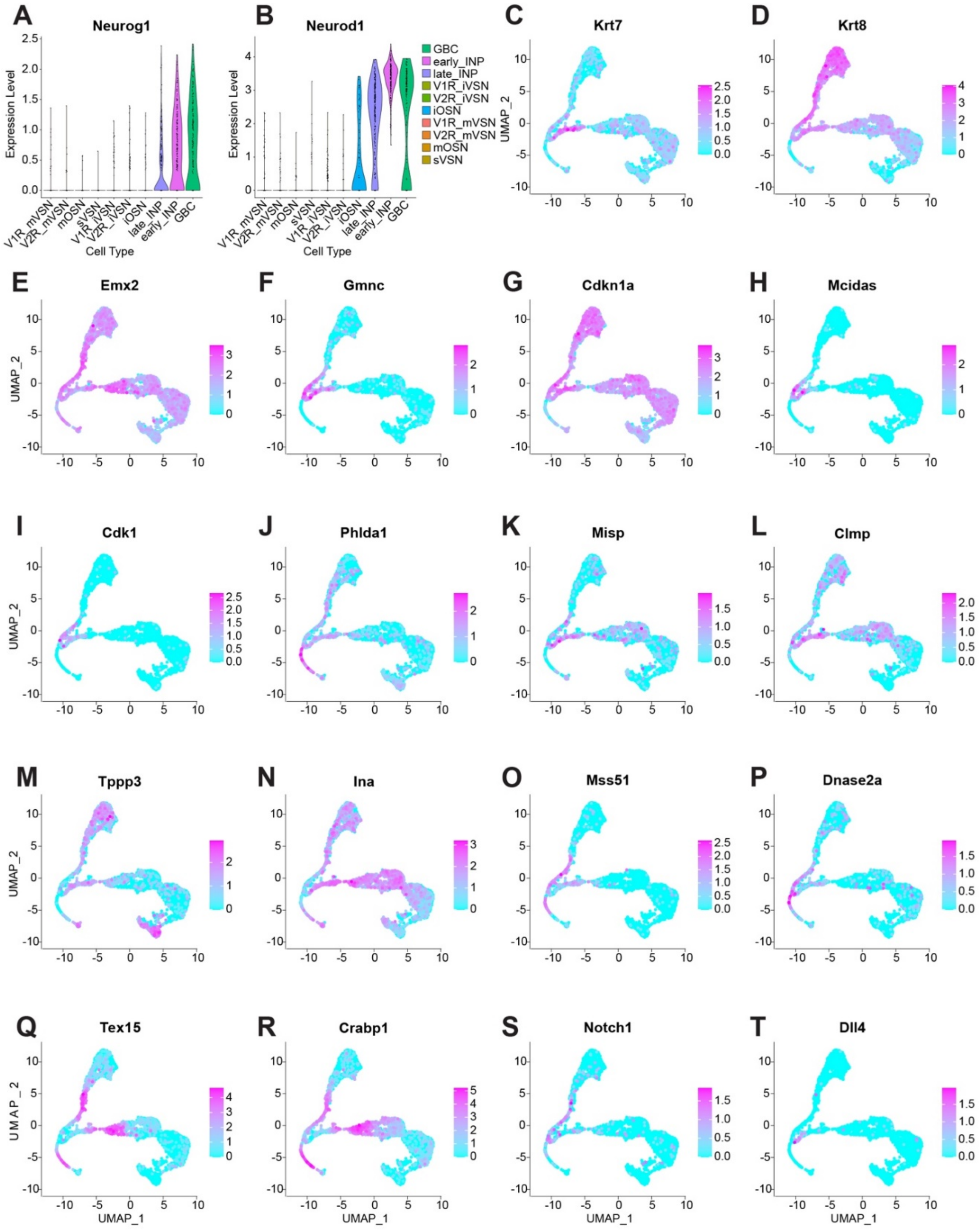


Fig. S7. Genes expressed during INP to immature neuron transition.

**A** and **B.** Violin plot of normalized expression for Neurog1 (A) and Neurod1 (B), split by cell type. **C**-**R**. Feature plots of normalized gene expression for early differentially expressed genes between V1R, V2R, and OSN lineages. **S** and **T**. Feature plots of normalized expression for Notch1 (S) and Dll4 (T).

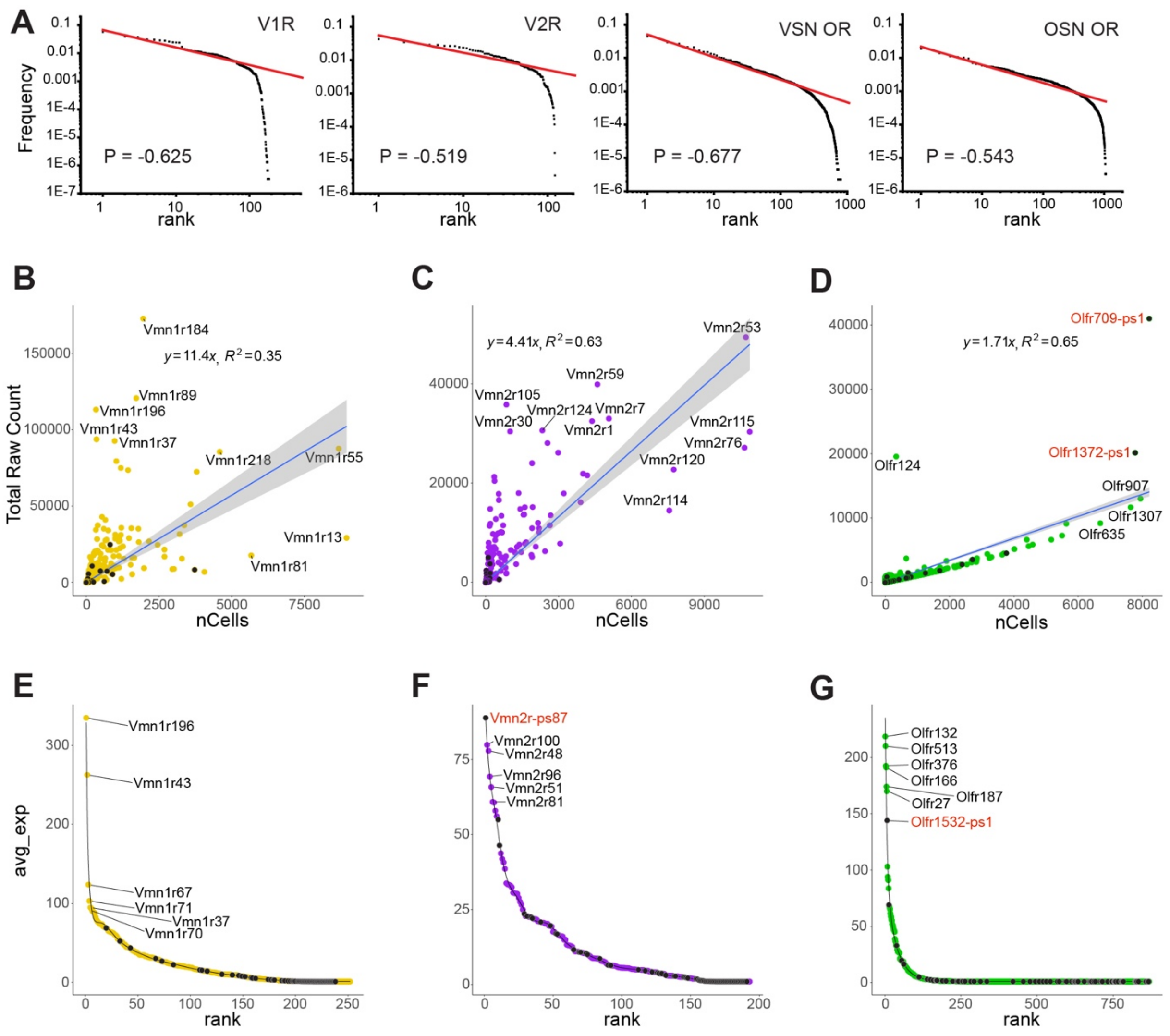


Fig. S8. Receptor distribution in the VNO.

**A**. Log scale rank-frequency distribution plots for V1R, V2R, VSN-OR, and OSN-OR. **B-D**. Number of cells expressing (nCells) vs. total raw counts for V1R, V2R, and VSN-OR. **E-F**. Rank by average count distributions for V1R, V2R, and VSN-OR.

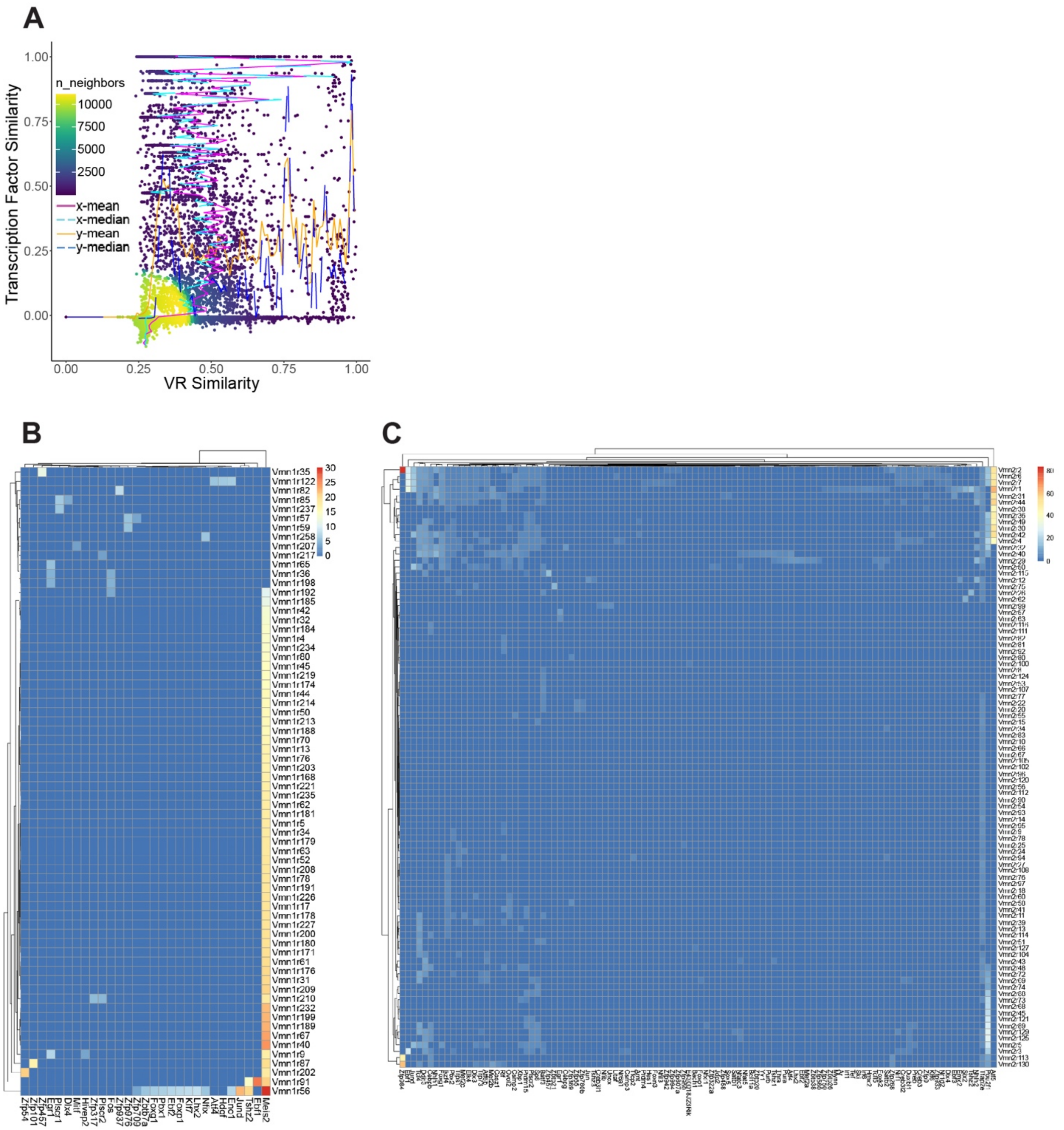


Fig. S9. Co-expression between vomeronasal receptors and transcription factors
**A**. Distribution density plot showing relationship between similarity of transcription factor (TF) gene expression profiles and receptor sequence similarity. **B-C**. Heatmaps showing the V1R-TF (B) and V2R-TF (C) associations. Heat shows average expression level for TF genes for a given receptor type.

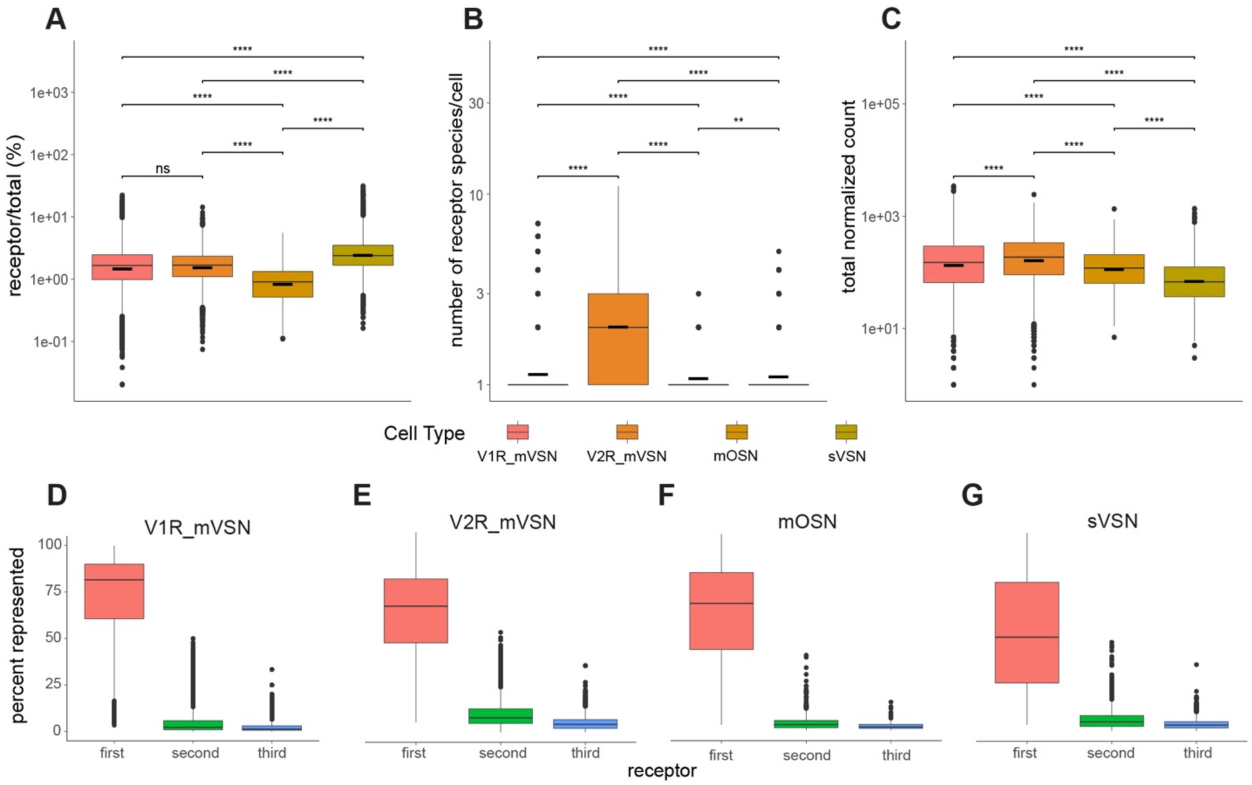


Fig. S10. Receptor expression statistics.
**A-C**. Percent of all read counts from a receptor gene (A), number of receptor species present per cell (B), and total counts from vomeronasal receptors (C), separated by cell type. **D-G**. Proportions of first, second, and third most expressed receptor gene as percent of total receptor counts, separated by cell type.
